# HRPK-1, a conserved KH-domain protein, modulates microRNA activity during *Caenorhabditis elegans* development

**DOI:** 10.1101/569905

**Authors:** Li Li, Isana Veksler-Lublinsky, Anna Y. Zinovyeva

## Abstract

microRNAs (miRNAs) are potent regulators of gene expression that function in diverse developmental and physiological processes. Argonaute proteins loaded with miRNAs form the miRNA Induced Silencing Complexes (miRISCs) that repress gene expression at the post-transcriptional level. miRISCs target genes through partial sequence complementarity between the miRNA and the target mRNA’s 3’ UTR. In addition to being targeted by miRNAs, these mRNAs are also extensively regulated by RNA-binding proteins (RBPs) through RNA processing, transport, stability, and translation regulation. While the degree to which RBPs and miRISCs functionally interact to regulate gene expression is likely extensive, we have only begun to unravel these functional interactions. An RNAi-based screen of putative ALG-1 Argonaute interactors has identified a role for a conserved RNA binding protein, HRPK-1, in modulating miRNA activity during *C. elegans* development. Here, we report the physical and genetic interaction between HRPK-1 and ALG-1/miRNAs. Specifically, we report the genetic and molecular characterizations of *hrpk-1* and its role in *C. elegans* development and miRNA-mediated target repression. We show that loss of *hrpk-1* causes numerous developmental defects and enhances the mutant phenotypes associated with reduction of miRNA activity, including those of *lsy-6, mir-35*-family, and *let-7*-family miRNAs. In addition to *hrpk-1* genetic interaction with these miRNA families, *hrpk-1* is required for efficient regulation of *lsy-6* target *cog-1*. We report that *hrpk-1* may play a role in miRNA processing but is not globally required for mature miRNA biogenesis or ALG-1/AIN-1 miRISC assembly and confirm HRPK-1 ability to co-precipitate with ALG-1. We suggest that HRPK-1 may functionally interact with miRNAs on multiple levels to enhance miRNA/miRISC gene regulatory activity and present several models for its activity.

**Author summary:** microRNAs are small non-coding RNAs that regulate gene expression at the post-transcriptional level. The core microRNA Induced Silencing Complex (miRISC), composed of Argonaute, mature microRNA, and GW182 protein effector, assembles on the target messenger RNA and inhibits translation or leads to messenger RNA degradation. RNA binding proteins interface with miRNA pathways on multiple levels to coordinate gene expression regulation. Here, we report identification and characterization of HRPK-1, a conserved RNA binding protein, as a physical and functional interactor of miRNAs. We confirm the physical interaction between HRPK-1, an hnRNPK homolog, and Argonaute ALG-1. We report characterizations of *hrpk-1* role in development and its functional interactions with multiple miRNA families. We suggest that HRPK-1 promotes miRNA activity on multiple levels in part by contributing to miRNA processing and by coordinating with miRISC at the level of target RNAs. This work contributes to our understanding of how RNA binding proteins and auxiliary miRNA cofactors may interface with miRNA pathways to modulate miRNA gene regulatory activity.

## Introduction

Robust regulation of gene expression is essential for normal development and cellular homeostasis. microRNAs (miRNAs), small non-coding RNAs ∼22nt in length, negatively regulate gene expression at the post-transcriptional level. miRNAs can act as developmental switches or can fine tune the expression of the target genes (for review, see [1], [2]). Processed miRNAs are loaded into their main protein cofactor, Argonaute (AGO), which then associates with members of the GW182 family of proteins, forming the microRNA Induced Silencing Complex (miRISC). Mature miRISCs bind to the target messenger RNAs (mRNAs) and repress their translation and/or destabilize the target mRNA [3], [4].

RNA binding proteins (RBPs) make up another class of post-transcriptional gene regulators. RBPs can affect miRNA gene-repressive activity in a variety of ways, including through miRNA processing [5] and mRNA co-targeting through mRNA processing, transport, localization, stability/degradation and mRNA translation regulation. A given mRNA bound by miRISC also serves as a platform for binding of additional RNA-interacting factors, and a few have been shown to associate with core miRISC and modulate its activity [6], [7], [8]. For example, NHL-2 and CGH-1 physically interact with miRISC and enhance the repression of miRNA target genes [7], a process regulated by casein kinase II [9]. Furthermore, many genes, including RNA binding proteins (RBPs), have been identified as genetic interactors of the *let-7* family of miRNAs [10], [11], [12], [13]. It remains unclear how many of them function as direct modulators of miRISC activity. Complementary to these approaches, we have previously sought to identify physical interactors of ALG-1, a miRNA-specific *C. elegans* Argonaute [14], hypothesizing that proteins that co-precipitate with ALG-1 include factors that modulate miRNA-induced gene repression. An RNAi-based screen of the putative ALG-1 co-factors has identified HRPK-1, a conserved homolog of human heterogeneous nuclear ribonucleoprotein K (hnRNPK), as a novel miRNA interactor that modulates activity of several miRNAs. Here, we report the characterization of both physical and genetic interactions between HRPK-1 and miRNA-containing complexes.

Heterogeneous nuclear ribonucleoproteins (hnRNPs) form a large family of nucleic acid binding proteins with diverse functions in a wide range of cellular processes, including transcription, RNA processing, RNA transport, RNA stability, and translational repression [15]. They possess multiple domains and are thought to form modular complexes, increasing the diversity of the RNA/protein interactions [16]. Several hnRNPs have recently emerged as having roles in miRNA-mediated gene regulation [15]. Some are implicated in biogenesis of specific miRNAs [17], while others, such as hnRNP Q, appear to hinder miRNA activity by competing with key molecular effectors of miRISC [18]. KH domain proteins are a subclass of hnRNPs with similarly diverse functions [15]. These proteins contain the KH nucleic acid binding motifs, which were originally identified in heterogeneous nuclear ribonucleoprotein K (hnRNPK), with KH domain named for <u>K</u> <u>h</u>omology [19]. KH domains are evolutionary conserved and are found in approximately 40 of human genes, and 27 of *C. elegans* genes. These nucleic acid binding motifs are 70 amino acids in size and can be present as a single domain or in multiple copies within a given protein [20], [19]. For example, Vigilin, a ribosome associated protein, contains 14 KH domains [21], [22]. It is thought that each KH domain can act as an independent binding module [23], although it is not currently clear if that holds true for all KH-containing proteins. Interestingly, a *C. elegans* homolog of the Vigilin gene, *vgln-1*, has been recently shown to cooperate with miRNAs to regulate gene expression [24].

Recent studies have implicated the human hnRNPK in miRNA-mediated gene repression. One study reports that hnRNPK may compete with multiple miRNAs for 3’UTR binding of their target gene, *PLK1* [25]. Such target binding competition puts hnRNPK in a role of a negative regulator of miRNA activity [25]. Other studies have placed hnRNPK in close physical proximity to miRNAs themselves, observing hnRNPK binding to miR-122 miRNA directly and potentially regulating its stability [26] or binding near miR-122 target sites on target mRNAs [27]. The functional significance of these interactions has not yet been described. In *C. elegans,* two KH domain-containing proteins, GLD-1 and VGLN-1 have been shown to genetically and/or physically interact with miRNA machinery [28] and [24], respectively. The mechanism through which GLD-1 and VGLN-1 functionally interact with the miRNA pathways remains unclear.

We have previously identified HRPK-1(F26B1.2), an hnRNPK homolog, in ALG-1 Argonaute immunoprecipitates as a putative ALG-1 physical interactor [14]. Here, we report the characterization of *hrpk-1* as a functional miRNA interactor that modulates activity of several miRNAs. Loss of *hrpk-1* results in a number of developmental defects, including sterility, embryonic lethality, vulval bursting, loss of alae, and abnormal gonad formation. Genetic loss of HRPK-1 enhances miRNA reduction-of-function phenotypes, consistent with HRPK-1 functional requirement for wild type miRNA activity. We report that HRPK-1 is ubiquitously expressed throughout *C. elegans* development and localizes to both nuclei and cytoplasm in some tissues, while remaining strongly nuclear in others. We confirm that HRPK-1 co-precipitates ALG-1 by immunoprecipitation. Finally, while *hrpk-1* appears to be globally dispensable for miRNA biogenesis, it may play a role in processing of select miRNAs. HRPK-1 is not needed for ALG-1/AIN-1 miRISC assembly, suggesting that HRPK-1 may also functions at the level of miRNA target repression, especially in case of miRNAs whose processing does not depend on HRPK-1 activity.

Our data suggest that HRPK-1 may regulate miRNA activity by interacting with miRNA-associated protein complexes and modulating the efficacy of both mature miRNA processing and miRNA activity on target mRNAs. This furthers our understanding of how miRNAs may be regulated by or function in concert with RNA binding proteins. Our findings also demonstrate that RNA binding proteins may integrate distinct developmental and physiological signals with miRNA-mediated gene repression.

## Results

To identify and characterize potential physical and functional interactions of HRPK-1 with miRISC components, we first generated a null allele in *hrpk-1.* A previously existing deletion allele of *hrpk-1, hprk-1(tm5522),* causes an in-frame deletion within the *hrpk-1* gene, resulting in production of a truncated HRPK-1 protein (Fig 1A-C). Using CRISPR/Cas9-based genome editing, we generated two independent null alleles of *hrpk-1, hrpk-1(zen15)* and *hrpk-1(zen17)* (Fig1A-C). Both alleles nearly completely or completely delete the *hrpk-1* locus (Fig 1A) and produce no HRPK-1 protein (Fig 1B-C). Since both null alleles produce a similar loss-of function phenotype (S1 Fig), we have designated the larger deletion, *hrpk-1(zen17)*, as the reference null allele for *hrpk-1* and used it in subsequent analyses.

**Fig 1.**
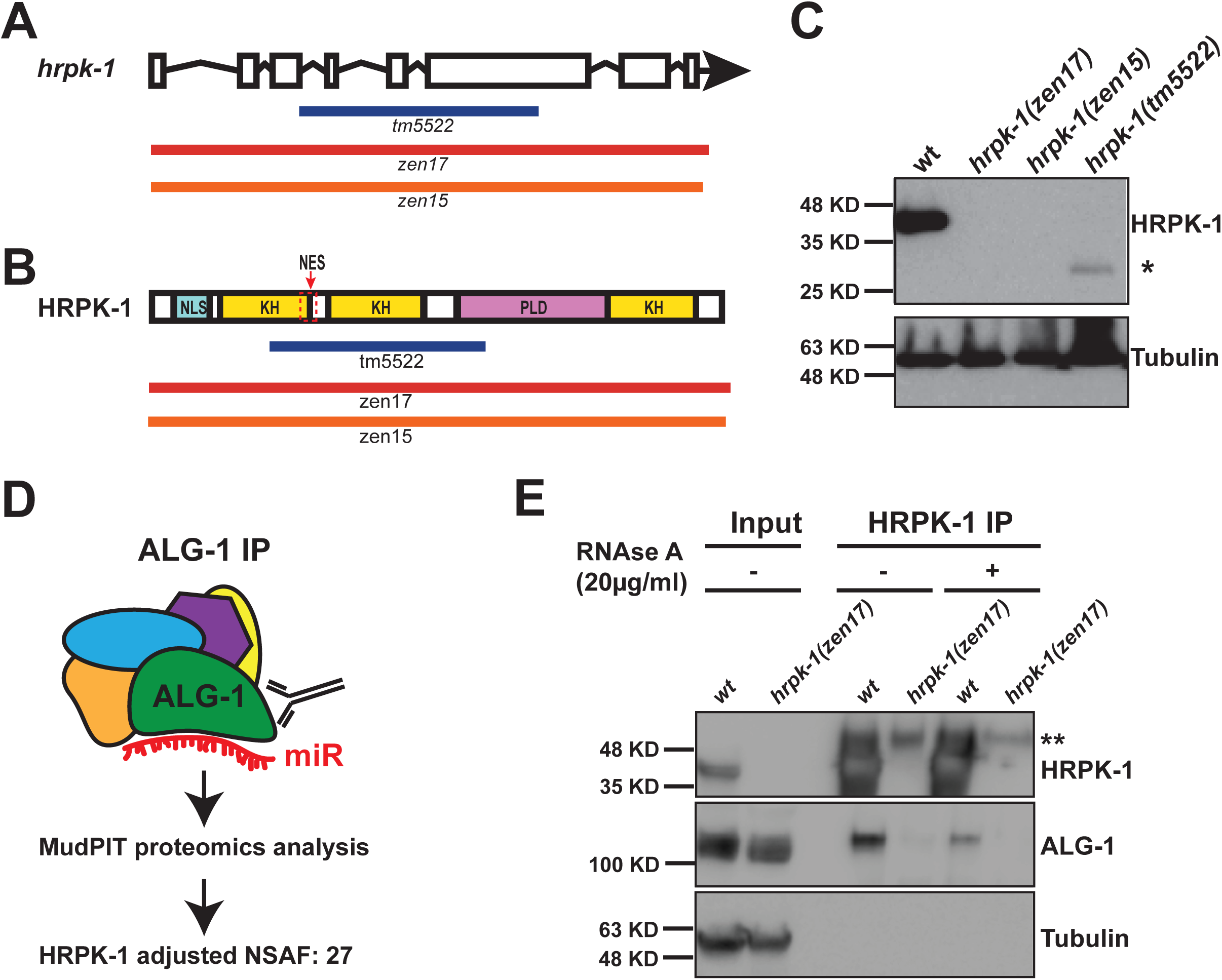
Multiple *hrpk-1* alleles produce deletions within the *hrpk-1* locus; ALG-1 co-precipitates with HRPK-1. (**A**) A schematic showing the predicted exon/intron structure of *hrpk-1* gene. *hrpk-1* alleles *tm5522, zen17,* and *zen15* delete parts or all of *hrpk-1*. (**B**) The *hrpk-1(tm5522)* allele is predicted to produce a truncated form of HRPK-1 protein, removing half of the first KH domain, second KH domain, and part of the PLD domain. NLS—predicted nuclear localization signal, KH—KH domain, NES—predicted nuclear export signal, PLD—prion like domain. (**C**) Western blotting for HRPK-1 protein detects the HRPK-1(tm5522) truncated protein at ∼27kDA (highlighted by *). No protein is detected from the *hrpk-1(zen17)* and *hrpk-1(zen15)* null animals. Tubulin is detected as a loading control. (**D**) HRPK-1 was identified by MudPIT mass spectrometry in ALG-1 co-precipitates [14]. NSAF = normalized spectral abundance factor. (**E**) Western blotting for HRPK-1 and ALG-1 proteins in the HRPK-1 immunoprecipitates treated and untreated with RNAse A. Tubulin is detected as a loading control for input. Input = 10% of IP. **this non-specific band is most likely antibody heavy chain.

### ALG-1 Argonaute and HRPK-1 co-immunoprecipitate in a partially RNA dependent manner

We have previously identified HRPK-1(F26B1.2) by immunoprecipitating *C. elegans* miRNA-specific Argonaute ALG-1 and analyzing the co-purified protein complexes by MudPIT proteomics [14], (Fig 1D). To confirm this physical interaction, we performed reciprocal IP using antisera generated against HRPK-1 and probed for the presence of ALG-1. We found that HRPK-1 IP co-precipitates ALG-1 (Fig 1E), consistent with HRPK-1 interaction with the miRNA machinery.

To determine whether HRPK-1/ALG-1 co-immunoprecipitation is RNA dependent, we performed the HRPK-1 IP in the presence of RNAse A (20μg/ml). Interestingly, incubation of lysates with RNAse A prior to and during HRPK-1 IP reduces but does not abolish ALG-1 co-precipitation with HRPK-1 (Fig 1E). This result suggests that while the ALG-1/HRPK-1 binding may in part depend on RNA, the two proteins may also be interacting directly, rendering the observed interaction partially RNAse A resistant (Fig 1E). The ALG-1/HRPK-1 interaction may be strengthened through RNA-protein interactions.

#### *hrpk-1* is required for a number of developmental processes

To determine what effects *hrpk-1* mutations have on animal development, we characterized gross morphological phenotypes of both *hrpk-1(tm5522)* and *hrpk-1(zen17)* mutants. We found that both alleles induce temperature sensitive sterility (Fig 2A), embryonic lethality (Fig 2B), and reduced brood size (Fig 2C). Both alleles also cause gonad formation defects (Fig 2D, E) and vulval bursting in day 3 or older adults (Fig 2F). We observed that almost in every instance, homozygous *hrpk-1(tm5522)* mutants exhibit more severe phenotypes than complete loss of *hrpk-1* (Fig 2A-C,G), with the exception of gonad morphology (Fig 2D,E) and vulval bursting (Fig 2F) defects. *hrpk-1(tm5522)* mutant animals also fail to produce adult alae approximately 40% of the time (Fig 2G). In addition, *hrpk-1* function appears to be maternally required for fertility (S2 Fig A), brood size (S2 Fig B), and embryonic viability (S2 Fig C). Specifically, *hrpk-1(zen17) [m-z-]* animals have increased defects compared to *hrpk-1(zen17) [m+z-]* in sterility (S2 Fig A), brood size (S2 Fig B), and embryonic lethality (S2 Fig C).

**Fig 2.**
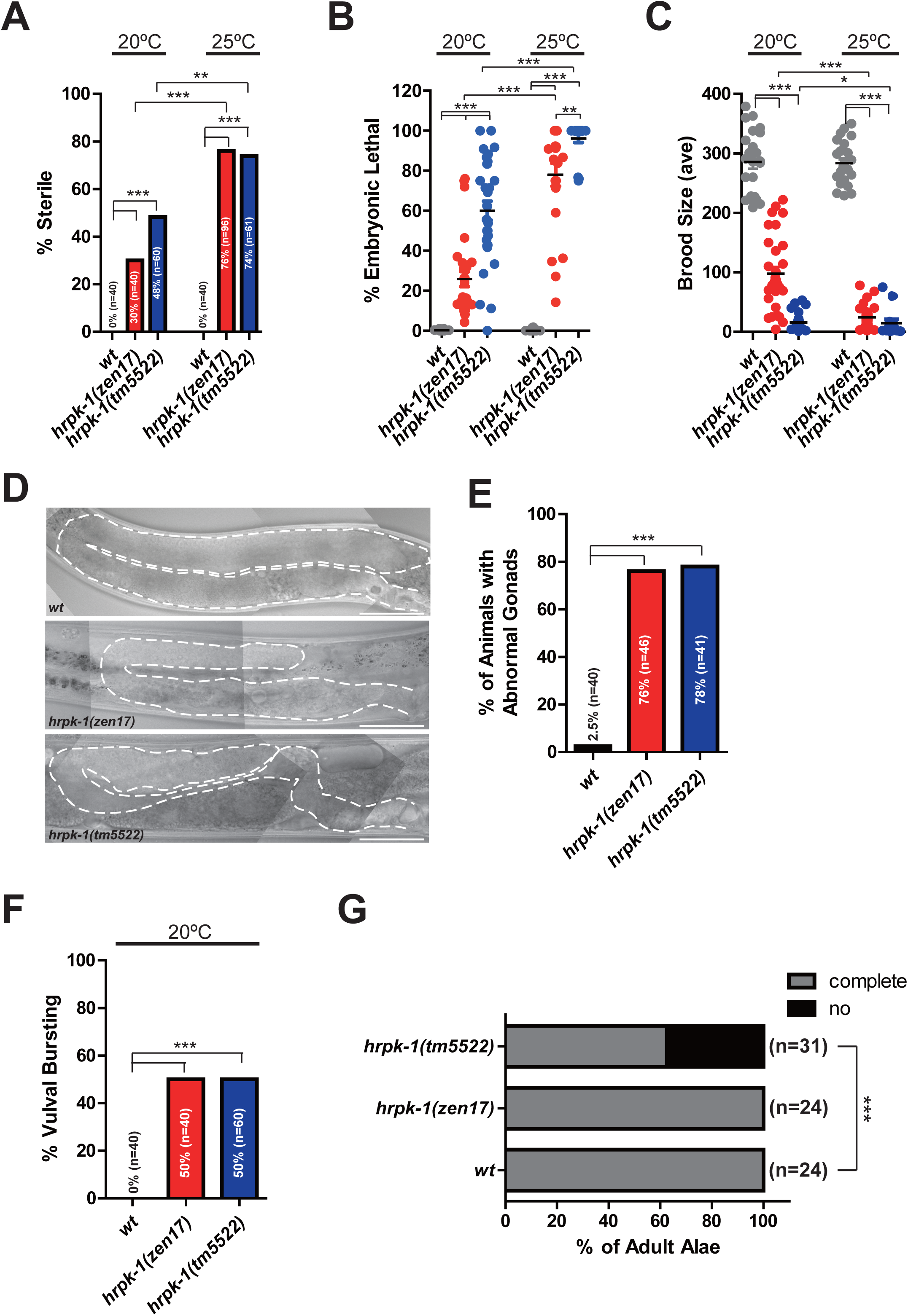
*hrpk-1* mutations cause developmental defects at 20°C and 25°C. *hrpk-1(zen17)* and *hrpk-1(tm5522)* mutations result in temperature sensitive animal sterility (**A**), embryonic lethality (**B**), and reduced brood size (**C**). *hrpk-1(zen17)* and *hrpk-1(tm5522)* mutations cause abnormal gonad formation (**D**, quantified in **E**), vulval bursting in day 3 or older adults (**F**), and defects in young adult alae formation (**G**). Bar in (**D**)=50μm. *\*p≤0.05, **p≤0.01, ***p≤0.001.*

#### *hrpk-1(tm5522)* allele is antimorphic

Our genetic analysis suggests that *hrpk-1(tm5522)* allele is weakly semi-dominant and antimorphic in nature (S2 Fig). Firstly, homozygous *hrpk-1(tm5522)* mutants almost always exhibit more severe phenotypes than *hrpk-1(zen17)* null animals (Fig 2A-C, G). Secondly, *hrpk-1(tm5522)/+ [m-z+]* animals carrying only a single copy of the wild type allele exhibited higher fertility defects and vulval bursting than *hrpk-1(zen17)/+ [m-z+]* animals (14% sterility versus 0%, respectively, S2 Fig A and D). Thirdly, *hrpk-1(tm5522)/+ [m+z+]* animals show a modest reduction in brood size (S2 Fig B) and embryonic lethality (S2 Fig C) and vulval bursting (S2 Fig D) not normally observed in wild type animals. Furthermore, animals carrying a single copy of maternally provided *hrpk-1(tm5522)* allele (*hrpk-1(tm5522) 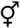 /hrpk-1(zen17)*♂ produced by *hrpk-1(tm5522)* mothers and *hrpk-1(zen17)* fathers) exhibit higher rates of sterility (S2 Fig A), have smaller brood sizes (S2 Fig B), and higher embryonic lethality (S2 Fig C) than *hrpk-1(zen17) [m-z-]* animals lacking functional *hrpk-1* completely. Animals carrying a single copy of maternally provided *hrpk-1(tm5522)* allele (*hrpk-1(tm5522) 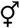 /hrpk-1(zen17)*♂also show more severe phenotypes than *hrpk-1(tm5522)*♂/ *hrpk-1(zen17) 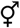* animals produced by *hrpk-1(tm5522)* fathers and *hrpk-1(zen17)* mothers (S2 Fig A-C), likely a result of both the antimorphic nature of the *tm5522* allele and the maternal component to its function. Overall, while the genetic analysis is somewhat complex due to the maternal requirement for *hrpk-1* activity, taken together, these genetic data support the antimorphic, weakly semi-dominant nature of the *hrpk-1(tm5522)* allele.

#### *hrpk-1* functionally interacts with the *let-7 family* of miRNAs

To test whether *hrpk-1* functions in miRNA-dependent cell fate specification, we examined potential genetic interactions between *hrpk-1* and *let-7* family of miRNAs that control temporal cell fate specification throughout larval development. The *let-7* miRNAs regulate the stage specific cell gene expression programs in a number of tissues, with the *let-7* family mutant phenotypes most evident in the *C. elegans* seam and hypodermis (for review, see [29]). Specifically, *let-7* family miRNAs, together with other heterochronic genes, regulate cell division patterns of the seam cells during larval development [30] and seam cell terminal differentiation at the transition from larval development to adulthood [31]. *mir-48, mir-241,* and *mir-84,* three members of the *let-7* miRNA family, are transcriptionally upregulated in the late L1 stage and subsequently down-regulate the expression of the *hbl-1* transcription factor during the L2 stage [30]. As a consequence of these regulatory interactions, *mir-48, mir-241,* and *mir-84* limit the proliferative seam cell division pattern of hypodermal stem cells to the L2 stage and promote subsequent L3-associated patterns of seam cell divisions [30], (Fig 3A). *mir-48, mir-241,* and *mir-84* are genetically redundant, with single and double mutants of these microRNAs exhibiting partially penetrant cell retarded heterochronic phenotypes [30]. These phenotypes can be monitored by observing alterations in seam cell lineage as well as defects associated with a reduction in the expression of adult-specific reporters (*col-19∷gfp*) after the 4^th^ larval stage [30,32], (Fig 3B). *hrpk-1(zen17)* enhances the retarded phenotype of *mir-48 mir-241 (nDf51)* mutants (Fig 3B-D). Specifically, loss of *hrpk-1* activity enhances the retarded expression of adult hypodermal marker, *col-19∷gfp(maIs105),* (Fig 3C and D) and results in an increased number of seam cells as compared to *mir-48 mir-241(nDf51)* alone (Fig 3B). This enhancement of the *mir-48 mir-241(nDf51)* heterochronic phenotype by loss of *hrpk-1* is consistent with *hrpk-1* functional requirement for efficient activity of the remaining intact *let-7* family miRNAs.

**Fig 3.**
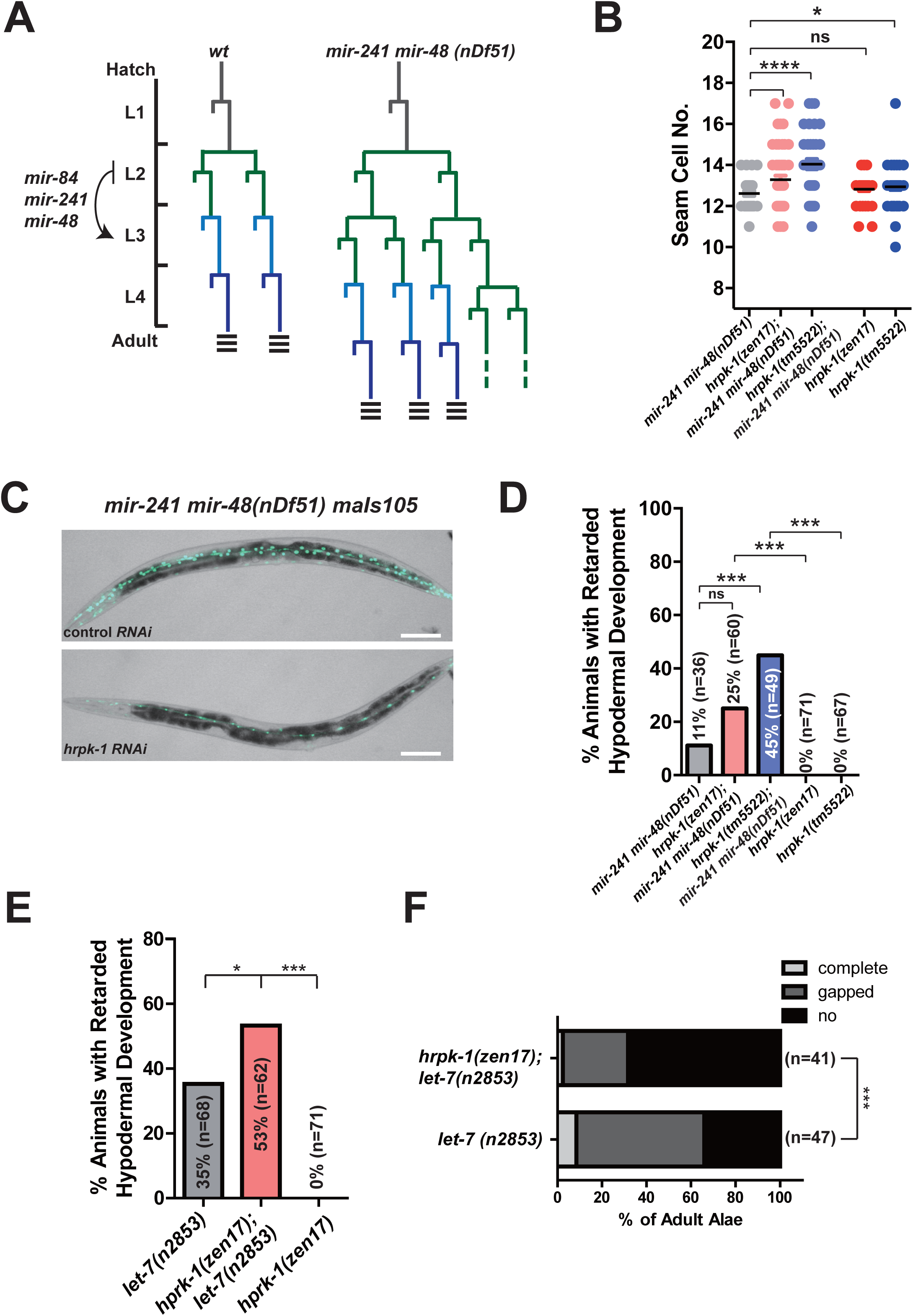
*hprk-1* functionally interacts with the *let-7* family of miRNAs. (**A**) Schematic seam cell lineages of wild type and *mir-241 mir-48* mutant animals throughout larval development. (**B**) *hrpk-1* mutations increase the seam cell numbers in *mir-48 mir-241(nDf51)* young adults. (**C**) Loss of wild type *hrpk-1* activity enhances the retarded hypodermal *col-19∷gfp* expression phenotype of *mir-48 mir-241(nDf51)* mutants (quantified in (**D**)). *hrpk-1(zen17)* enhances the retarded (**E**) alae formation phenotype and (**F**) *col-19∷gfp* expression observed in *let-7(n2853)* animals at 20°C. Animals scored in (**B**-**E**) carry the *col-19∷gfp(maIS105)* transgene. All animals in (**E, F**) carry *lin-2(e1309),* which suppresses the vulval bursting of *let-7(n2853)* animals through a non-heterochronic mechanism. Bar in (**C**)=100μm. \**p≤0.05, **p≤0.01, *** p≤0.001, **** p≤0.0001.*

*let-7,* the founding member of the *let-7* family of miRNAs, controls terminal cell fate specification, which occurs during the developmental progression through the late larval stages and into the adulthood [31]. A temperature sensitive mutation that compromises but does not completely abolish let-7 activity, *let-7(n2853)*, causes a retarded development phenotype [31], [33]. *let-7(n2853)* animals have abnormal/absent alae in young adults and delayed hypodermal expression of the adult marker *col-19∷gfp* at 20°C (Fig 3E and F), [31]. *hrpk-1* knockout enhances retarded *col-19∷gfp(maIS105)* expression (Fig 3E) and the retarded alae phenotype (Fig 3F) observed in the *let-7(n2853)* mutants. Loss of *hrpk-1* alone is not enough to induce a heterochronic phenotype (Fig 3A, D, C). The observed enhancement of the *let-7(n2853)* retarded phenotype by *hrpk-1* mutations is consistent with the hypothesis that *hrpk-1* function is important for *let-7* miRNA activity.

#### *hrpk-1* functionally interacts with the *lsy-6* miRNA and is required for efficient regulation of *lsy-6* target *cog-1*

To test the hypothesis that HRPK-1 may be required for miRNA activity regulation, we looked for the effects of *hrpk-1* mutations on the activity of *lsy-6* miRNA-dependent processes. *lsy-6* regulates cell fates of two bilaterally symmetrical ASE neurons [34]. ASEL-specific expression of *lsy-6* down-regulates its key target, *cog-1*, while uninhibited *cog-1* expression within the ASER dictates that neuron’s cell fate [34]. The ASEL cell fate is distinguished by the expression of a downstream reporter, *Plim-6∷gfp,* an established marker for the ASEL cell fate [34], [35], (Fig 4A). Reduction of lsy-6 activity through a cis-regulatory mutation in the *lsy-6* promoter, *lsy-6(ot150),* causes a low penetrance phenotype where the ASEL neuron adopts the cell fate of ASER approximately 20% of the time [36], (Fig 4A and B). To assess *hrpk-1-lsy-6* functional interaction we removed *hrpk-1* in the presence of the reduction of function *lsy-6(ot150)* allele. Genetic mutations in *hrpk-1* significantly enhance the cell fate defective phenotype observed in the *lsy-6(ot150)* animals (Fig 4B). *hrpk-1* mutations alone are not sufficient to induce an ASEL to ASER cell fate switch (Fig 4B). Importantly, *hrpk-1* RNAi relieves the lsy-6-mediated inhibition of its target, *cog-1∷gfp,* in uterine cells (Fig 4C-D), suggesting that HPRK-1 is required for efficient inhibition of *cog-1* by lsy-6 miRNA in that tissue.

**Fig 4.**
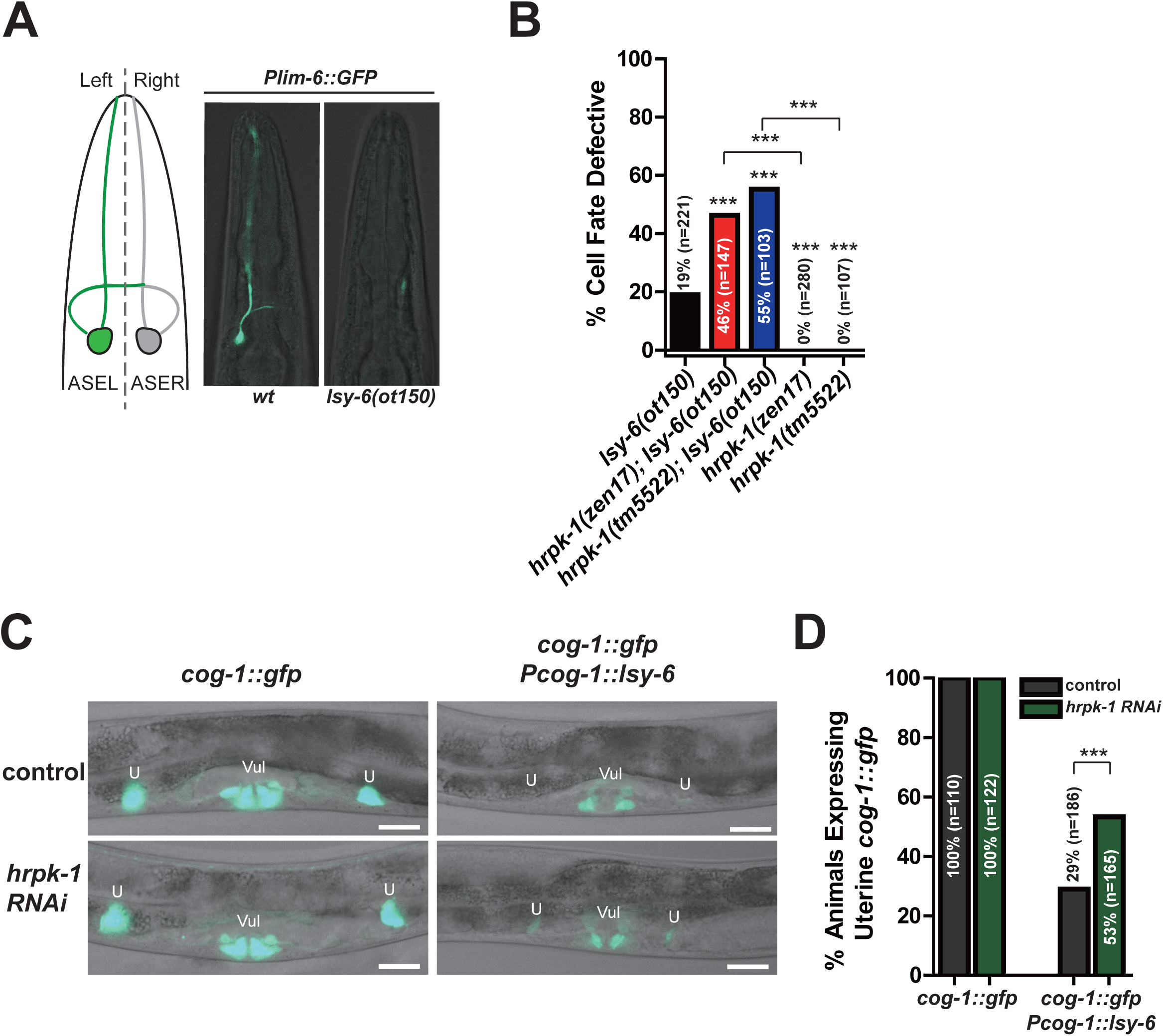
*hrpk-1* function is required for efficient *lsy-6* miRNA activity. *Plim-6∷gfp* reporter marks the ASEL cell fate (**A**). The low penetrance defective cell fate phenotype observed in *lsy-6(ot150)* animals is enhanced by *hrpk-1* mutations (**B**). (**C**) *hrpk-1* RNAi relieves the lsy-6-mediated repression of *cog-1∷gfp* in the uterine cells, quantified in (**D**). Animals in (B) also carry *Plim-6∷gfp* reporter. Bar in (C)=20μm. *\*\*\*p<0.001.*

#### *hrpk-1* functionally interacts with the *mir-35-42* family of miRNAs

To assess whether *hrpk-1* may be broadly required for miRNA activity, we reduced its function in other miRNA sensitized backgrounds. *mir-35-42* family of miRNAs controls fertility and embryonic development of *C. elegans* [37]. Loss of the entire *mir-35-42* miRNA family results in fully penetrant sterility and embryonic lethality phenotypes [37]. Deletion of a genomic region harboring 7 of the 8 *mir-35* family miRNAs (*mir-35-41*), *mir-35-41(nDf50),* causes animals to exhibit temperature sensitive increase in embryonic lethality and overall reduction of brood size [37], [38], (Fig 5 A and B). Combining either *hrpk-1* deletions (*hrpk-1(zen17)* and *hrpk-1(tm5522))* with the *mir-35-41(nDf50)* deletion allele results in a strong synthetic lethal phenotype where a majority of animals fail to develop (Fig 5A). This genetic synergy in combined mutants is also recapitulated in a dramatic decrease in overall brood size (Fig 5B). In both cases, animals were reared at the semi-permissive temperature of 20°C. *hrpk-1(tm5522)* causes a greater enhancement of the *mir-35-41(nDf50)* phenotype than *hrpk-1(zen17)* null (Fig 5 A-B), consistent with the antimorphic nature of the *hrpk-1(tm5522)* allele. Importantly, the enhancement of the *mir-35-41(nDf50)* phenotype by *hrpk-1* mutations is not simply additive, but synergistic (Fig 5A-B), consistent with the hypothesis that *hrpk-1* activity may be required for remaining *miR-42* function.

**Fig 5.**
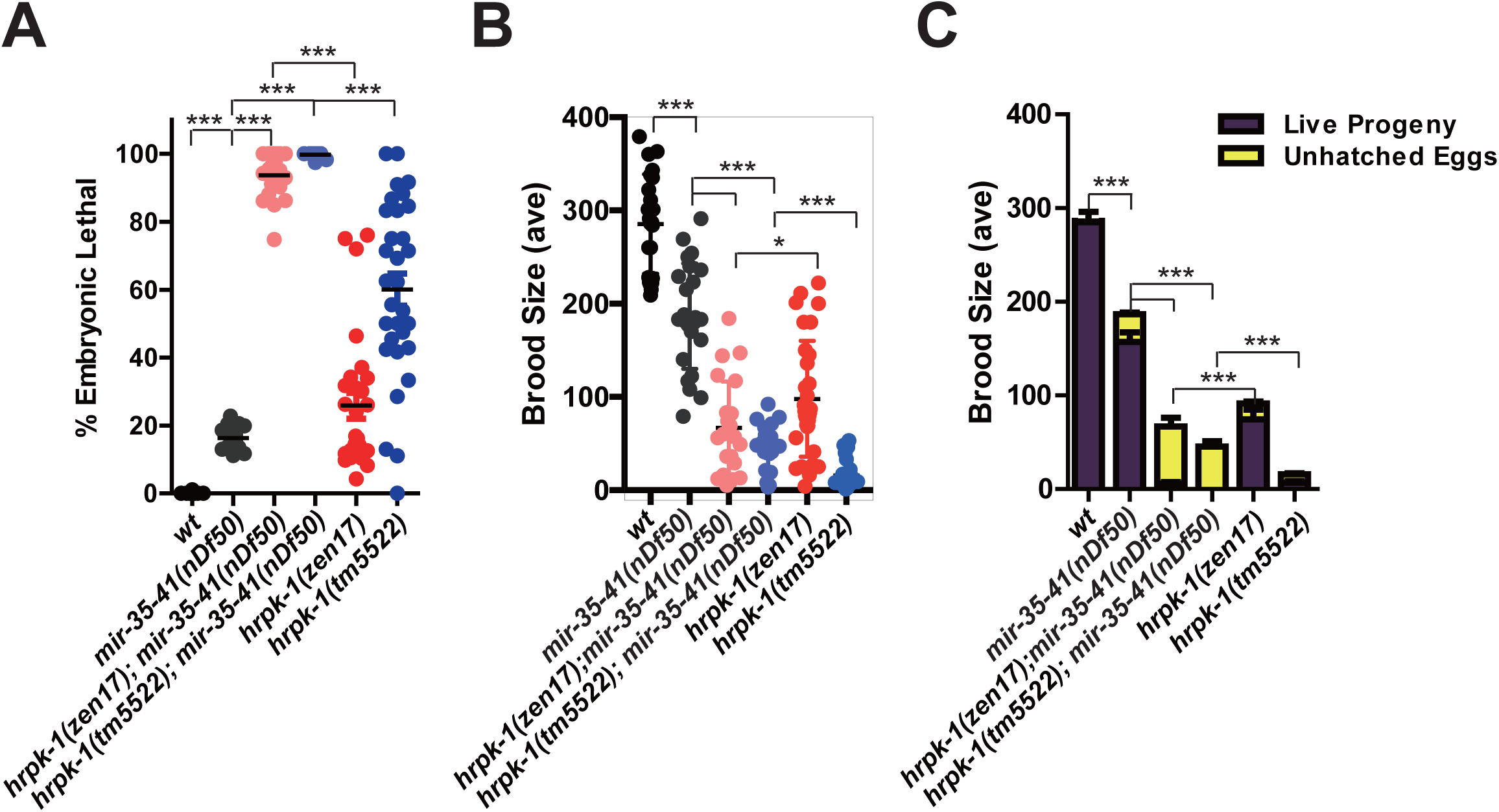
*hrpk-1* mutations enhance *mir-35-41(nDf51)* mutant phenotype at 20°C. *hprk-1* mutations enhance *mir-35-41(nDf51)* embryonic lethality (**A**) and further reduces *mir-35-41(nDf51)* brood sizes (**B-C**). (**C**) shows the brood breakdown by live and dead progeny. *\*p<0.05* and *\*\*\*p<0.001.*

### HRPK-1 is ubiquitously expressed throughout *C. elegans* development

To gain insight into HRPK-1 function, we characterized both spatial and temporal expression of endogenous HRPK-1 using a CRISPR-generated *hrpk-1∷gfp* transgene (Fig 6). *hrpk-1∷gfp* is ubiquitously expressed during all *C. elegans* developmental stages (Fig 6A-D). The C-terminally tagged *hrpk-1∷gfp* expression is observed in the gut, muscle, neuronal, and hypodermal tissues, where it localizes to the cell nuclei (Fig 6A, B). Interestingly, *hrpk-1∷gfp* is strongly present in the animal germline, oocytes, and early embryos, where its subcellular localization is both nuclear and cytoplasmic (Fig 6B-D). The subcellular localization of HRPK-1∷GFP is consistent with the predicted nuclear localization and nuclear export signals found within the *hrpk-1* sequence (Fig 1B). Our inability to detect *hrpk-1∷gfp* signal in the cytoplasm of somatic cells may reflect HRPK-1 distinct functions between the soma and the germline or may simply be due to a lower expression level that falls below our detection limits. In fact, HRPK-1 somatic cytoplasmic localization has been observed large scale subcellular proteome mapping [39], suggesting that we may be limited in detecting HRPK-1 in the cytoplasm in the presence of strong nuclear expression. Overall, the spatial and temporal *hrpk-1* pattern of expression is consistent with the apparent *hrpk-1* roles in a number of developmental processes, including embryonic and larval development, as well as fertility.

**Fig 6.**
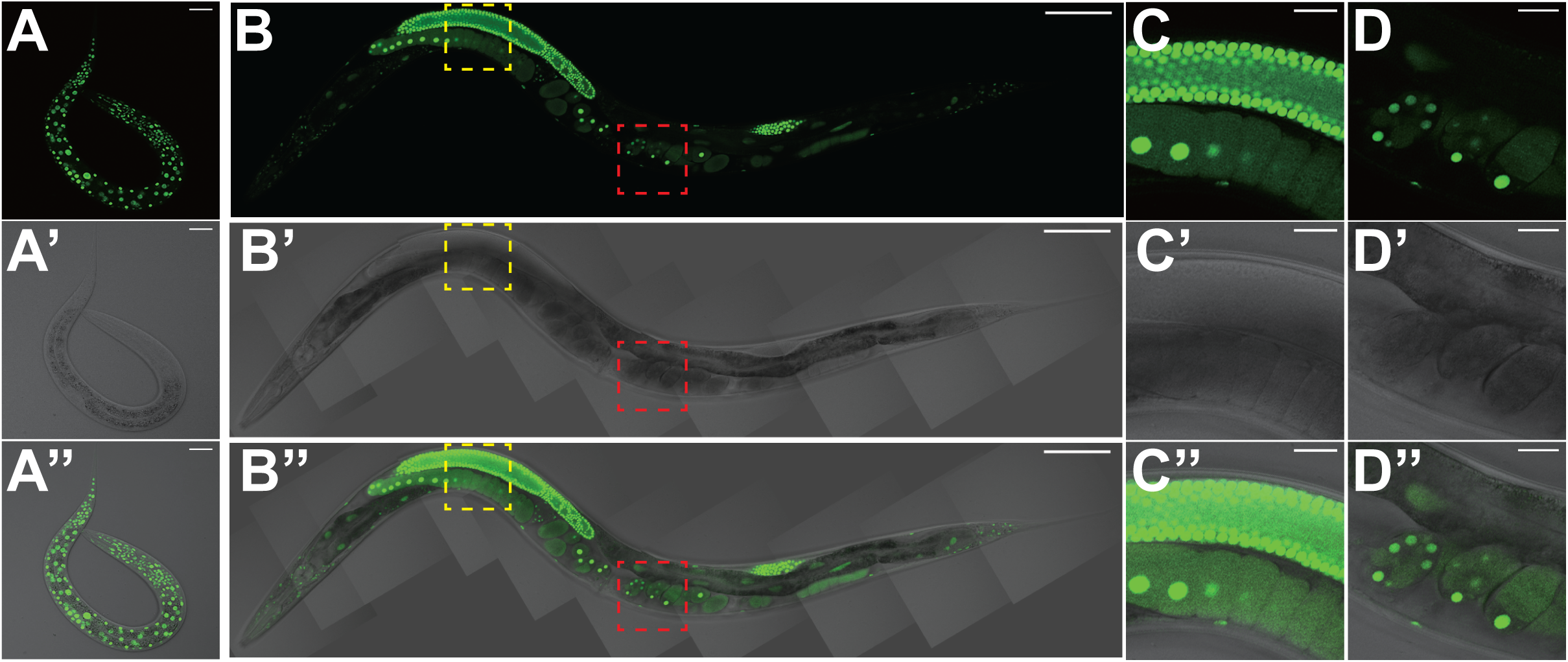
Endogenously tagged *hrpk-1∷gfp* transgene *(hrpk-1(zen64))* is expressed throughout *C. elegans* development. (**A**) L1 larva shows ubiquitous HRPK-1∷GFP expression in the nuclei of somatic tissues and the germline precursor cells. (**B**) HRPK-1∷GFP fluorescence in an adult animal highlights ubiquitous HRPK-1 expression in somatic tissues and in the animal’s germline. Zoom in micrographs of (**B**) highlight strong nuclear and cytoplasmic expression in the germline (**C**) and early embryos (**D**). Bar in (**A**), (**C**), (**D**)=20μm, bar in (**B**)=100μm.

### Loss of *hrpk-1* modestly affect levels of select miRNAs but does not affect ALG-1/AIN-1 miRISC assembly

Functional regulation of the miRNA activity by HRPK-1 could occur at the level of miRNA processing, miRISC assembly, or miRISC activity. To determine whether HRPK-1 is necessary for miRNA biogenesis, we cloned and sequenced miRNAs from wild type, *hrpk-1(zen17),* and *hrpk-1(tm5522)* mutant animals. We found that in general, abundance of mature miRNAs is not globally affected in *hrpk-1(zen17)* animals (Fig 7A-C, S1 Table), with only five mature miRNAs reduced 2 or more-fold in *hprk-1(zen17)* mutant and three miRNAs increased more than 2-fold (S2 Table). In contrast, *hrpk-1(tm5522)* animals have an expanded set of miRNAs reduced in abundance. Twenty-seven miRNAs, including members of the mir-35 family, were reduced 2-fold or more in the *hrpk-1(tm5522)* animals, and seven miRNAs were increased in abundance in *htpk-1(tmm5522)* mutants (S2 Table). We did observe a >2-fold decrease in lsy-6 abundance in both *hrpk-1(zen17)* and *hrpk-1(tm5522)* animals (Fig 7D and S2 Table). This reduction in lsy-6 counts could explain the functional interaction observed between *hrpk-1(-)* and *lsy-6(ot150)*. In contrast, let-7, mir-84, and mir-42 levels were not consistently decreased in *hrpk-1* mutants (Fig 7D and S1 Table). In fact, mir-84 counts were somewhat increased in both *hrpk-1(zen17)* and *hrpk-1(tm5522)* animals and let-7 and mir-42 were slightly decreased in the *hrpk-1(tm5522)* mutants only (Fig 7D and S1 Table). These data suggest that the functional interaction between *hrpk-1* mutations and *let-7(n2852), mir-48 mir-241(nDf51),* and *mir-35-41(nDf50)* miRNA mutants does not arise from a decrease in miRNA abundance of the remaining intact miRNAs (let-7, mir-84, and mir-42).

**Fig 7.**
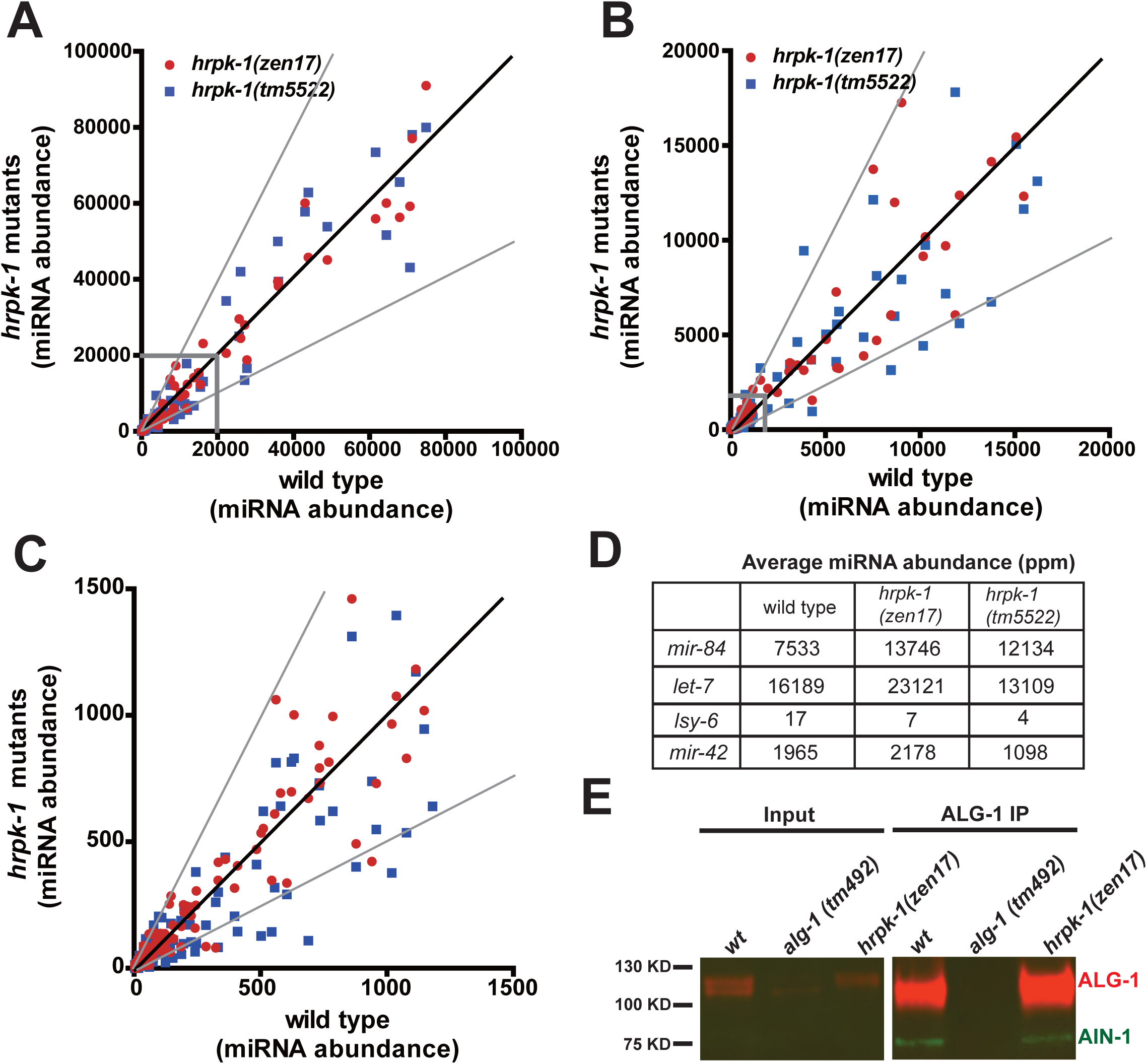
miRNA abundances and miRISC formation in *hrpk-1* mutant animals. Mature miRNA abundance in wild type and *hrpk-1* mutant animals (**A-C**). The black line represents a perfect 1:1 correlation between wild type and *hrpk-1* mutant miRNA abundances. Grey lines delineate a 2-fold difference in abundance. (**B**) Zoom in of grey box in (**A**). (**C**) Zoom in of the grey box in (**B**). Both miRNA and miRNA* abundances are graphed. (**D**) Average miRNA abundances for miRNAs whose function is affected by *hrpk-1* mutations in our functional assays. (**E**) AIN-1 co-precipitates with ALG-1 in *hrpk-1* mutant animals to levels similar to wild type.

Mature miRISC formation involves Argonaute association with GW182 homologs [4]. To assess whether HRPK-1 may be needed for miRISC assembly, we tested the ability of ALG-1 to co-precipitate with its miRISC co-factor and GW182 homolog AIN-1 in *hrpk-1(zen17).* We found that AIN-1 co-precipitates with ALG-1 in *hrpk-1* mutant animals at levels similar to those observed in wild type (Fig 7E). We did not test for ALG-2 and AIN-2 containing miRISC formation and therefore cannot rule out the possibility that HRPK-1 may be necessary for formation of miRISCs that include ALG-2 and/or AIN-2 proteins. However, we can conclude that HRPK-1 is not required for ALG-1/AIN-1 interaction as observed by our ALG-1 immunoprecipitation experiments (Fig 7E).

## Discussion

### HRPK-1 physically and functionally interacts with miRNA pathways

In this manuscript we report characterizations of *hrpk-1,* which encodes an RNA binding protein HRPK-1. HRPK-1 was originally identified by MudPIT proteomics in ALG-1 co-precipitates [14], (Fig 1D). Here, we confirm that HRPK-1 can co-precipitate ALG-1 in a reciprocal experiment and show that the interaction between HRPK-1 and ALG-1 is only partially RNA-dependent (Fig1E). The relatively small amount of ALG-1 co-precipitating with HRPK-1 could reflect instability of complexes in our assay or may represent the portion of ALG-1 in complex with HRPK-1 (Fig 1E). Importantly, not only is HRPK-1 a conserved protein and an ortholog of human hnRNP K, the interaction between Argonaute and HRPK-1 also appears to be conserved as hnRNP K has previously been co-purified with three of four human AGO proteins [40].

In addition to confirming the conserved physical interaction between HRPK-1 and ALG-1, we report that *hrpk-1* is genetically required for efficient activity of several miRNAs and miRNA families: *lsy-6, let-7-family,* and *mir-35-family*. Loss of *hrpk-1* alone does not alter ASEL/ASER cell fate decisions despite enhancing the *lsy-6(ot150)* reduction-of-function phenotype. In addition to enhancing the cell fate defects observed in *lsy-6(ot150)* animals, HRPK-1 promoted lsy-6 mediated repression of lsy-6 target gene, *cog-1* (Fig 4C and D). Similarly, loss of *hrpk-1* did not cause a heterochronic phenotype normally observed in *let-7* family mutants (Fig 3), although the antimorphic *hprk-1(tm5522)* allele did produce a mild alae defect (Fig 2G). Together, these data suggest that HRPK-1 is not an essential miRNA co-factor, but rather positively modulates *lsy-6* and *let-7* family miRNA activity. The synergistic genetic interaction between *hrpk-1* mutations and *mir-35-41(nDf50)* is also consistent with a hypothesized role for *hrpk-1* as a positive co-factor of mir-42 activity. Overall, we show that HRPK-1 physically interacts ALG-1, a major component of miRLC/miRISC and functionally interacts with several miRNAs in a number of developmental processes.

#### *hrpk-1* subcellular localization

Overall, the ubiquitous expression of *hrpk-1* is consistent with its pleotropic effects on *C. elegans* development. We observed a strong nuclear expression of endogenous *hrpk-1* (Fig 6) and both nuclear and cytoplasmic HRPK-1∷GFP localization in the germline, oocytes, and early embryos. This is not surprising as HRPK-1 does contain a predicted nuclear localization signal and a predicted nuclear export signal (Fig 1B). Despite our inability to definitively detect HRPK-1∷GFP in the cytoplasm of somatic tissues, a recent report mapping the subcellular-specific proteomes in a number of *C. elegans* tissues places HRPK-1 in the hypodermal cytoplasm in addition to the nucleus using a proximity ligation dependent assay [39]. This assay may be able to capture transient protein shuttling and likely has a higher sensitivity than our fluorescent reporter observations. We would argue that the observed steady-state nuclear HRPK-1∷GFP localization does not mean there is no function for HRPK-1 in the cytoplasm. In fact, given the biochemical interaction between ALG-1 and HRPK-1, we hypothesize that cytoplasmic HRPK-1 activity may in fact be required for its functional interaction with the miRNA pathways. It remains to be seen whether nuclear HRPK-1 localization or movement between the nucleus and the cytoplasm plays a role in miRNA-dependent gene expression regulation.

### How might HRPK-1 enhance miRNA gene repressive activity?

Overall, several molecular models can explain the genetic and physical interactions between HRPK-1 and miRNA machinery. It is possible that HRPK-1 may interact with ALG-1 at the miRNA Loading Complex (miRLC) and participate in mature miRNA biogenesis (Fig 8A). We observed miRNA abundance changes for a small subset of miRNAs, including lsy-6, in *hrpk-1* mutant animals (Fig 7A-D and S2 Table). The observed functional interaction between *hrpk-1* and *lsy-6* mutations could therefore be explained by a reduced abundance of lsy-6 miRNA, potentially placing HRPK-1 in the *lsy-6* processing pathway. However, the abundances of the majority of miRNAs, including the *let-7-family* and *mir-42* were not consistently decreased between wild type and *hprk-1* mutants (S1 Table and Fig 7A-D). This observation suggests that the functional interaction between *hrpk-1* mutants and *let-7(n2853), mir-48 mir-241(nDf51),* and *mir-35-41(nDf50)* (Figs 3 and 5) is likely not due to *hrpk-1(-)* effects on miRNA processing. In fact, most mature miRNA abundances are not affected in *hrpk-1(zen17)* animals (Fig 7A-C, S1 Table). HRPK-1 involvement at the level of miRLC may therefore be limited to a small subset of miRNAs, including lsy-6. However, we cannot at this point rule out the possibility that HRPK-1 may play a broader role in miRLC formation or miRNA processing. The greater reduction of miRNA numbers in *hrpk-1(tm5522)* compared to *hrpk-1(zen17)* mutants hints at a possible explanation for the more severe phenotypes observed in *hrpk-1(tm5522)* animals, potentially through sequestration of components essential for miRNA processing or stability. Further work will be needed to characterize the extent to which HRPK-1 may be involved in miRNA processing and/or stability and the cause of the antimorphic nature of the *hrpk-1(tm5522)* allele.

**Fig 8.**
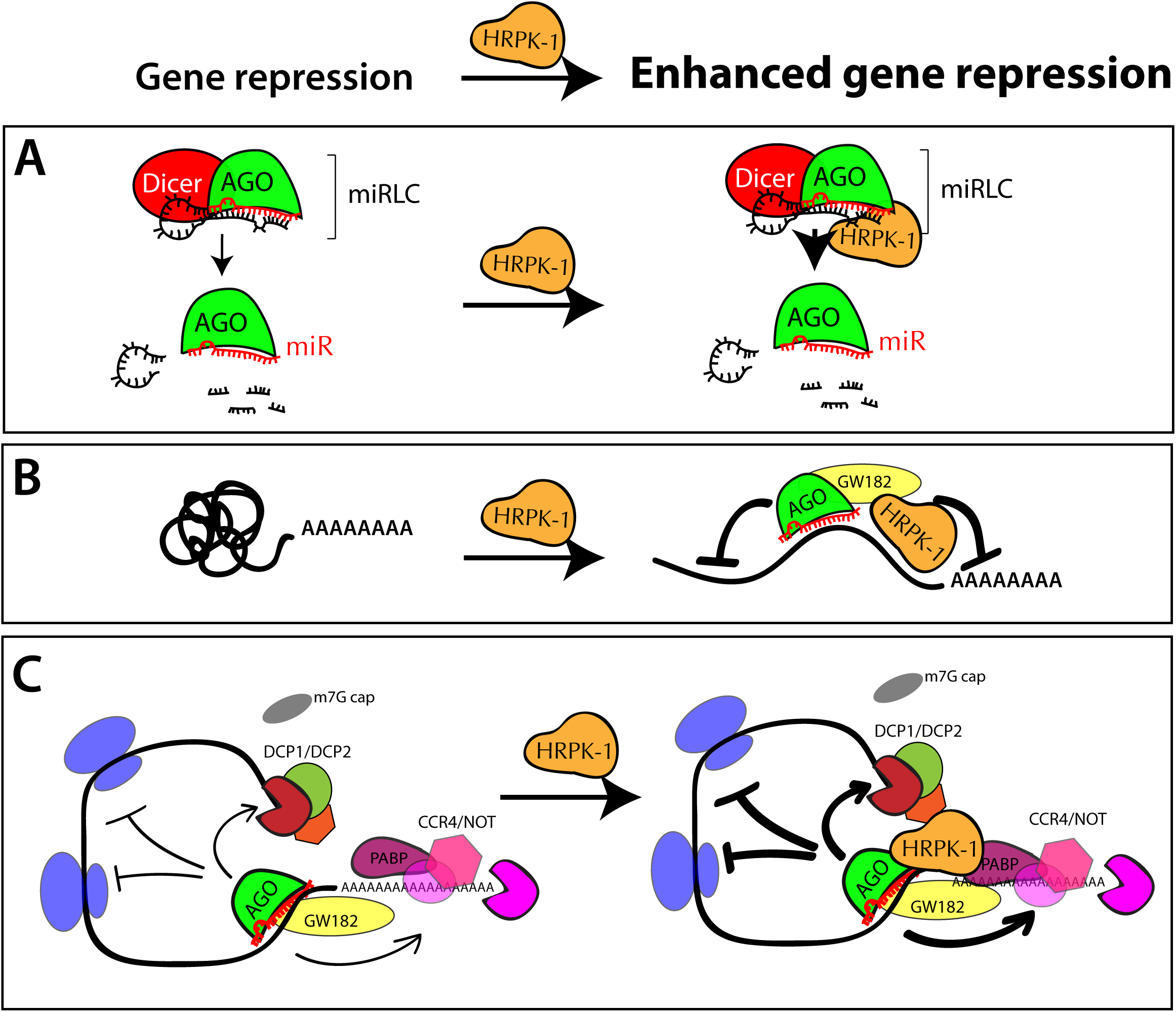
Models representing potential mechanisms through which HRPK-1 synergizes with the miRISC on mRNA targets. These models are not mutually exclusive. (**A**) HRPK-1 may cooperate with miRLC and assist in miRNA processing. (**B**) HRPK-1 may enhance miRISC activity by increasing miRNA target site availability through mRNA folding and enhancing miRISC∷RNA binding. (**C**) HRPK-1 may bind mRNAs at sites near miRNA target sites and enhance miRISC∷mRNA target interaction through miRISC binding. In this model, HRPK-1 may also increase miRISC activity through augmenting miRISC interaction with downstream effector complexes.

We found that HRPK-1 is not required for ALG-1/AIN-1 miRISC assembly, as *hrpk-1(zen17)* animals have an intact ALG-1/AIN-1 miRISC (Fig 7E). This observation is consistent with the hypothesis that HRPK-1 may interact with ALG-1 downstream of miRISC formation, possibly via the target mRNAs. We favor this hypothesis, as it is difficult to imagine a sole role for HRPK-1 at the level of miRLC that does not globally affect mature miRNA biogenesis or mature miRISC formation. Interestingly, the interaction between HRPK-1 and ALG-1 is only partially RNA-dependent (Fig 1E), suggesting that RNA is not necessary for this interaction to occur but may enhance it. HRPK-1 is a member of the KH domain family of proteins, which are known to be RNA binding proteins with diverse functions. These functions could include roles in mRNA processing, mRNA stability and folding, and mRNA transport out of the nucleus, and translational control. It is possible that, similar to other hnRNP proteins, HRPK-1 has diverse functions and may coordinate with miRNA pathways on multiple levels, perhaps even by bridging the miRLC with miRISC. In addition to a potential role in miRNA processing for subset of miRNAs (Fig 8A), HRPK-1 may stimulate miRNA/miRISC activity by increasing miRNA target site availability through mRNA folding (Fig 8B). In this model, HRPK-1 association with target mRNAs and miRISC may enhance miRISC accessibility and binding to target mRNAs and therefore increase the target gene repression (Fig 8B). In addition, HRPK-1 could stimulate the interaction between miRISC and downstream effector complexes (Fig 8C). Identification of HRPK-1 RNA binding sites and their proximity to miRNA target sites will in the future help characterize the physical relationship between the two.

Given the diversity of hnRNP and KH domain proteins functions, we must allow for the possibility that multiple, miRISC-dependent and independent HRPK-1 activity synergizes with the miRNA-mediated target regulation. For example, an HRPK-1 role in mRNA processing could increase miRNA site availability. Identifying the impacts of HRPK-1 loss on alternative splicing of mRNAs and the resulting changes in miRNA binding site availability can in the future shed light on the plausibility of this model. It should be noted that none of the proposed models are mutually exclusive and multiple HRPK-1 functions could synergize, directly and/or indirectly, with the miRNA gene regulatory pathways. We look forward to further investigating the mechanisms through which HRPK-1 and miRNAs cooperate to facilitate gene repression.

Overall, our results demonstrate that HRPK-1, a KH domain RNA binding protein, physically and functionally interacts with miRNA-mediated gene repression. Other KH domain RNA binding proteins have been shown to biochemically and genetically interact with the miRNA machinery. These include the *C. elegans* translational repressor, a Quaking homolog, and a KH domain protein, GLD-1, among others [28], [41] and VGLN-1 [24]. Additional KH-domain proteins and other hnRNPs have been detected in large-scale mass spectrometry experiments [41], [14], [42]. The molecular mechanisms by which RNA binding proteins synergize with miRNA pathways to regulate mRNA expression are diverse and remain largely unknown. Further studies to understand the mechanism of functional interactions between RBPs, including HRPK-1, and miRNA-mediated gene repression will shed light on how these two classes of post-transcriptional gene regulators cooperate to regulate animal development.

## Materials and methods

### Strain maintenance, RNAi, and phenotypic assessments

*C. elegans* strains were grown in standard conditions on NGM plates using OP50 as a food source [43]. Strains were maintained at 20°C unless otherwise noted. RNAi gene knockdown was performed as previously described [44]. Briefly, PS3662 or OH7310 animals were placed on *hrpk-1* RNAi food and the F1 progeny were scored for the *cog-1∷gfp* presence in vulval and uterine cells.

For the *mir-48 mir-241(nDf51)* assay, young adults were scored for seam and hypodermal *col-19∷gfp* expression, alae formation, and seam cell number. Seam cells numbers were obtained by counting the seam cells located between the pharynx and the anus of a given animal. Phenotypes were scored on either the Zeiss Axioplan 2 or the Leica DM6 upright microscopes equipped with DIC and epifluorescence. For *mir-35-41(nDf50)* assay, animals were scored for sterility, brood size and embryonic lethality. Animal sterility, embryonic lethality, and brood size were assessed by scoring entire broods of individual animals. Animals were considered sterile if they were unable to produce any progeny, dead or alive. Brood size refers to the total number of progeny produced, including dead embryos. Percent embryonic lethality was calculated as follows: # of dead embryos/total progeny (live progeny + dead embryos). Vulval bursting was visually assessed beginning at the Day 1 young adult stage and daily thereafter.

### CRISPR-based genome editing

*hrpk-1(zen15)* and *hrpk-1(zen-17)* deletion alleles were produced using CRISPR/Cas9 genome editing using *hrpk-1* site specific guides 1 and 9, corresponding to the 5’ and 3’ ends of the gene, respectively. The following *hrpk-1* guides were used to generate the deletions: *hrpk-1* guide 1, ACTGTCTGTTCCATTAATAG and *hrpk-1* guide 9, CAAGGCCGTGAACGATTCGG. *hrpk-1(zen17)* was outcrossed seven times and *hrpk-1(zen15)* was outcrossed twice. The entire *hrpk-1* locus was sequenced to confirm the nature of the deletion in each mutant.

Endogenously tagged *hrpk-1∷gfp* strains were generated by inserting a worm codon-optimized GFP coding sequence and a self-excising cassette (SEC) into the C-terminus of endogenous *hrpk-1* locus just before the stop codon through CRISPR/Cas-9 triggered homologous recombination [45]. The SEC was then excised as described [45]. The donor sequence was generated by subcloning 596 bp upstream of the *hrpk-1* stop codon and 600 bp downstream of *hrpk-1* stop codon into the pDD282 vector [45] using Hi Fi assembly kit (NEB). The *hrpk-1* stop codon was eliminated from the donor sequence to allow in frame GFP tag addition. A PAM site mutation corresponding to guide 9 was included in the donor DNA sequence resulting in the sequence change of [CAAGGCCGTGAACGATTCGG<u>TGG</u>] change to [CAAGGCCGTGAACGATTCGG<u>TCG</u>] immediately upstream of the GFP tag sequence. Animal microinjections were performed as previously described [46]. Six independent *hrpk-1∷gfp* lines were obtained and the resulting endogenously tagged *hrpk-1∷gfp* loci was sequenced. Since all six lines looked superficially wild type and showed the same *hrpk-1∷gfp* expression pattern, a single line was chosen for an in-depth analysis. The *hrpk-1∷gfp(zen64)* strain was outcrossed twice and assessed for *hrpk-1* functional integrity. *hrpk-1∷gfp* transgenic line had wild type fertility and embryonic viability (S3 Fig), confirming that the GFP-tagged HRPK-1 retained its wild type activity.

The following primers were used in generation of the *hrpk-1* donor sequence in order to place the GFP sequence at the C-terminus of *hrpk-1*:

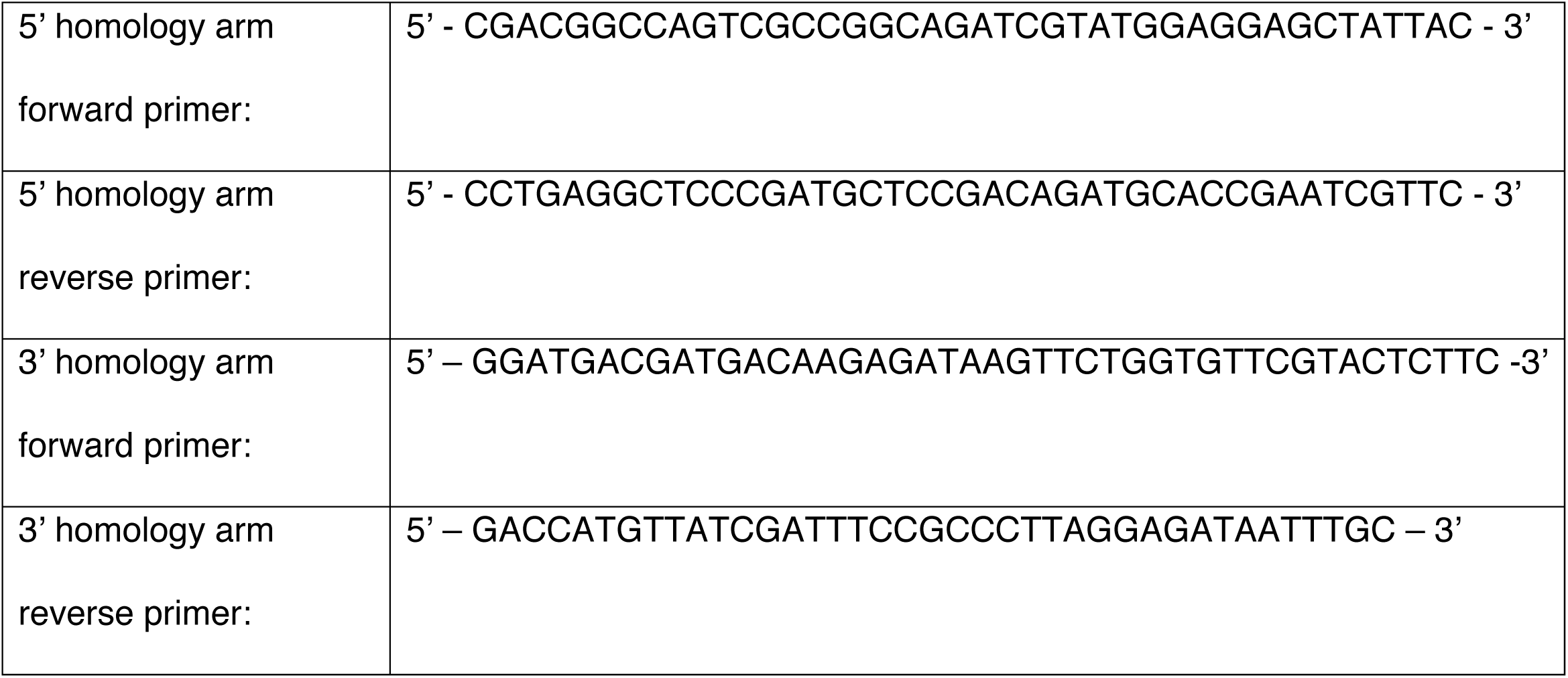

### Extract preparation and immunoprecipitation (IP)

Worm extracts were prepared as previously described [47] with the following modifications. Mixed stage animals were homogenized using a Bullet blender (MIDSCI). Briefly, 300*μ*L worm pellets were mixed with RNAse free Rhino beads (MIDSCI) and 300*μ*L lysis buffer and homogenized at the highest setting for 4 minutes. Homogenate was then moved to a fresh tube and spun at 13,000xg for 20 minutes to clarify the extract. The extracts were then used for immunoprecipitation experiments or were flash frozen in liquid nitrogen and stored at −80°C.

ALG-1 immunoprecipitation and Western blotting were performed as previously described [48] using Protein A Dynabeads (ThermoFisher). HRPK-1 detection was done using a custom rabbit anti-HRPK-1 antibody (Pocono) generated against the C-terminal peptide of HRPK-1 (CVRNSTQGRERFGGSV) at the 1:1000 dilution. HRPK-1 immunoprecipitation was performed using the same method as ALG-1 immunoprecipitation [48], with the anti-HRPK-1 antibody covalently crosslinked to the Protein A Dynabeads (ThermoFisher) using dimethylpimelimidate1. Mouse antitubulin antibody (Sigma-Aldrich) was used to detect tubulin as a loading control.

### RNA preparation and small RNA sequencing

RNA from mixed stage animals was prepared as previously described [47] with the following modifications. Mixed stages animals resuspended in 250μL water were mixed with 1ml of Trizol (Fisher Scientific) and RNAse free Rhino beads (MIDSCI) and homogenized using a Bullet blender (MIDSCI) at the highest setting for 4 minutes. Homogenate was then moved into a fresh tube, mixed with 212μL of chloroform, and spun down to separate the phases. All remaining steps of the RNA purification were performed as previously described [47].

For small RNA libraries preparation, small RNAs were first size selected by gel purification as described in [49]. The size selected RNA was then used to construct small RNA libraries using the NEXTflex Small RNA Library Prep kit v3 (Bioo Scientific) and sequenced on the Illumina HighSeq instrument at the Kansas University Genome Sequencing Core. Data analysis was performed as previously described [14]. The total mapped reads across two replicates were as follows: N2 (7,386,035), UY38 (7,679,177), and UY42 (7,540,761).

### Microscopy and statistics

Localization of the endogenously tagged HRPK-1∷GFP transgene was imaged using Zeiss Axioplan 2 upright microscope equipped with a Zeiss Axiocam HR digital camera. Images were assembled using Photoshop. Contrast and brightness of images were not adjusted. F-test was used to analyze brood size, t-test was used to analyze embryonic lethality, and chi-square was used to analyze all other phenotypic data.

### Strains used in this study

*N2 wild type*

*UY44 hrpk-1(zen15)*

*UY38 hrpk-1(zen17)*

*UY39 hrpk-1(zen17); maIs105[col-19∷gfp]*

*UY42 hrpk-1(tm5522)*

*UY43 hrpk-1(tm5522); maIs105[col-19∷gfp]*

*OH3646 otls114[Plim-6∷gfp + rol-6(su1006)]; lsy-6(ot150)*

*UY46 hrpk-1(zen17) otIs114[Plim-6∷gfp + rol-6(su1006)]; lsy-6(ot150)*

*UY54 hrpk-1(zen17) otIs114[Plim-6∷gfp + rol-6(su1006)]*

*UY9 hrpk-1(tm5522) otIs114[Plim-6∷gfp + rol-6(su1006)]; lsy-6 (ot150)*

*UY18 hrpk-1(tm5522) otls114[Plim-6∷gfp + rol-6(su1006)]*

*PS3662 syIs63[cog-1∷gfp + unc-119(+)]*

*OH7310 otIs193 [cog-1p∷lsy-6hp + rol-6(su1006)] syIS63[cog-1∷gfp + unc-119(+)]*

*MT14119 mir-35-41(nDf50)*

*UY56 hrpk-1(zen17)/ht2gfp; mir-35-41(nDf50)*

*UY55 hrpk-1(tm5522)/ht2gfp; mir-35-41(nDf50)*

*VT1296 mir-241,mir-48(nDf51) maIs105[col19∷GFP]*

*UY67 hrpk-1(zen17); mir-241 mir-48(nDf51) maIs105[col19∷gfp]*

*UY68 hrpk-1(tm5522); mir-241 mir-48(nDf51) maIs105[col19∷gfp]*

*HML11 maIs105[col19∷gfp] lin-2 (e1309) let-7 (n2853)*

*UY157 hrpk-1(zen17); maIs105[col19∷gfp] lin-2 (e1309) let-7 (n2853)*

*VT3841 alg-1(tm492)*

*UY66 hrpk-1∷gfp(zen64)*

## Acknowledgements

We are grateful to the Caenorhabditis Genetics Center (CGC), funded by NIH Office of Research Infrastructure Programs (P40 OD010440), which provided some of the strains used in this project. We thank the National BioResource Project (NBRP) at the Tokyo Women’s Medical University School of Medicine for providing the *hrpk-1(tm5522)* allele described in this study. We thank Yin Wang for technical support. We are grateful to the Ambros lab, where the HRPK-1 interaction with ALG-1 was initially discovered. We thank C. Hammell for strains and the critical reading of this manuscript, and M. Han for AIN-1 antisera. This work has been supported by Kansas INBRE, P20GM103418 to L.L. and A.Y.Z., and R35GM124828 to A.Y.Z.

---

**S1 Fig.**
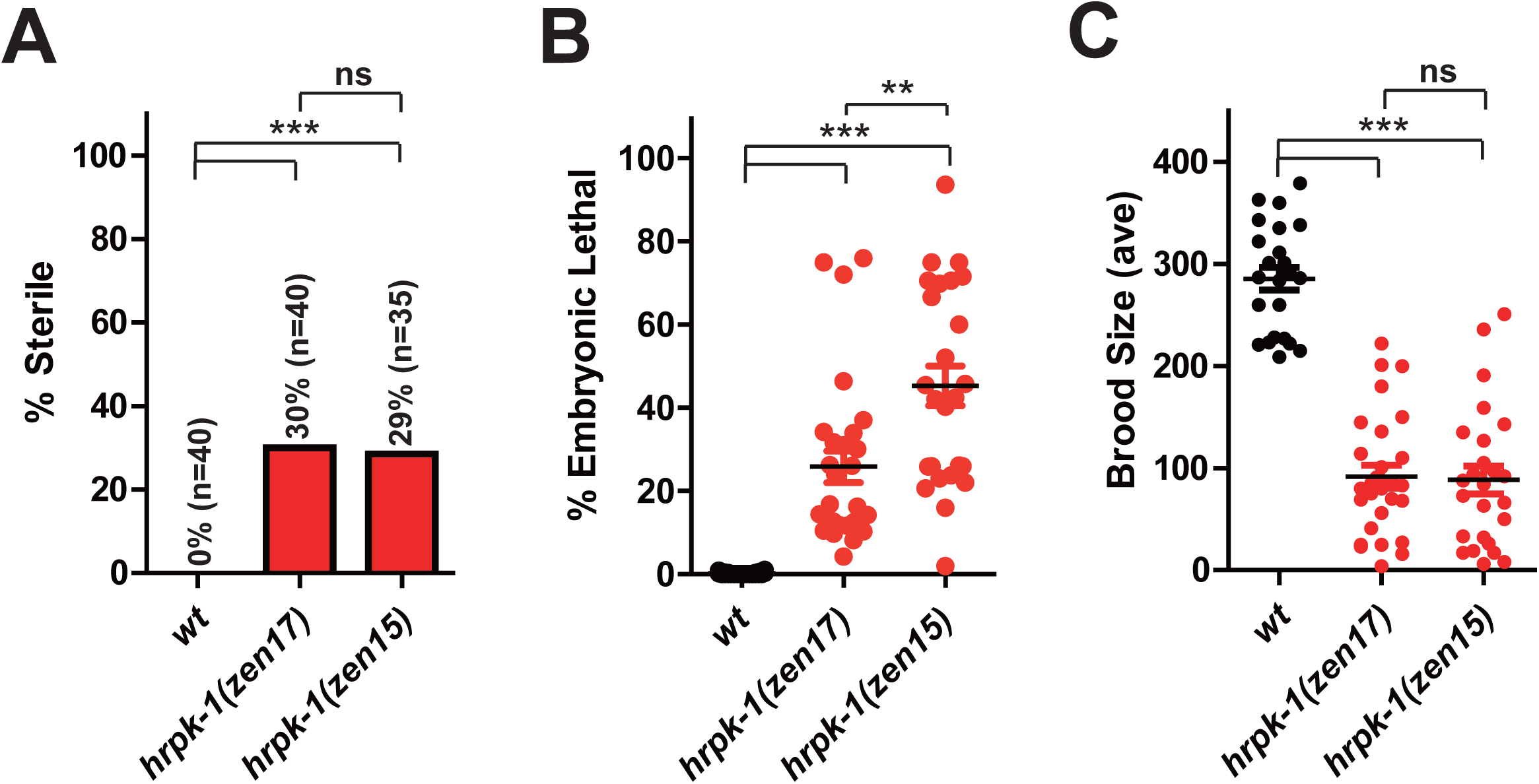
*hrpk-1(zen17)* and *hrpk-1(zen15)* show similar phenotypes. *hrkp-1(zen15)* and *hrpk-1(zen17)* produce similar levels of sterility (**A**), embryonic lethality (**B**), and brood size (**C**). An increase in embryonic lethality in *hrpk-1(zen15)* mutant animals (**D**) is most likely due the difference in the number of outcrosses between the strains, with *hrpk-1(zen17)* being outcrossed seven times, while *hrpk-1(zen15)* was outcrossed only twice. \*\*\**p≤0.001*

**S2 Fig.**
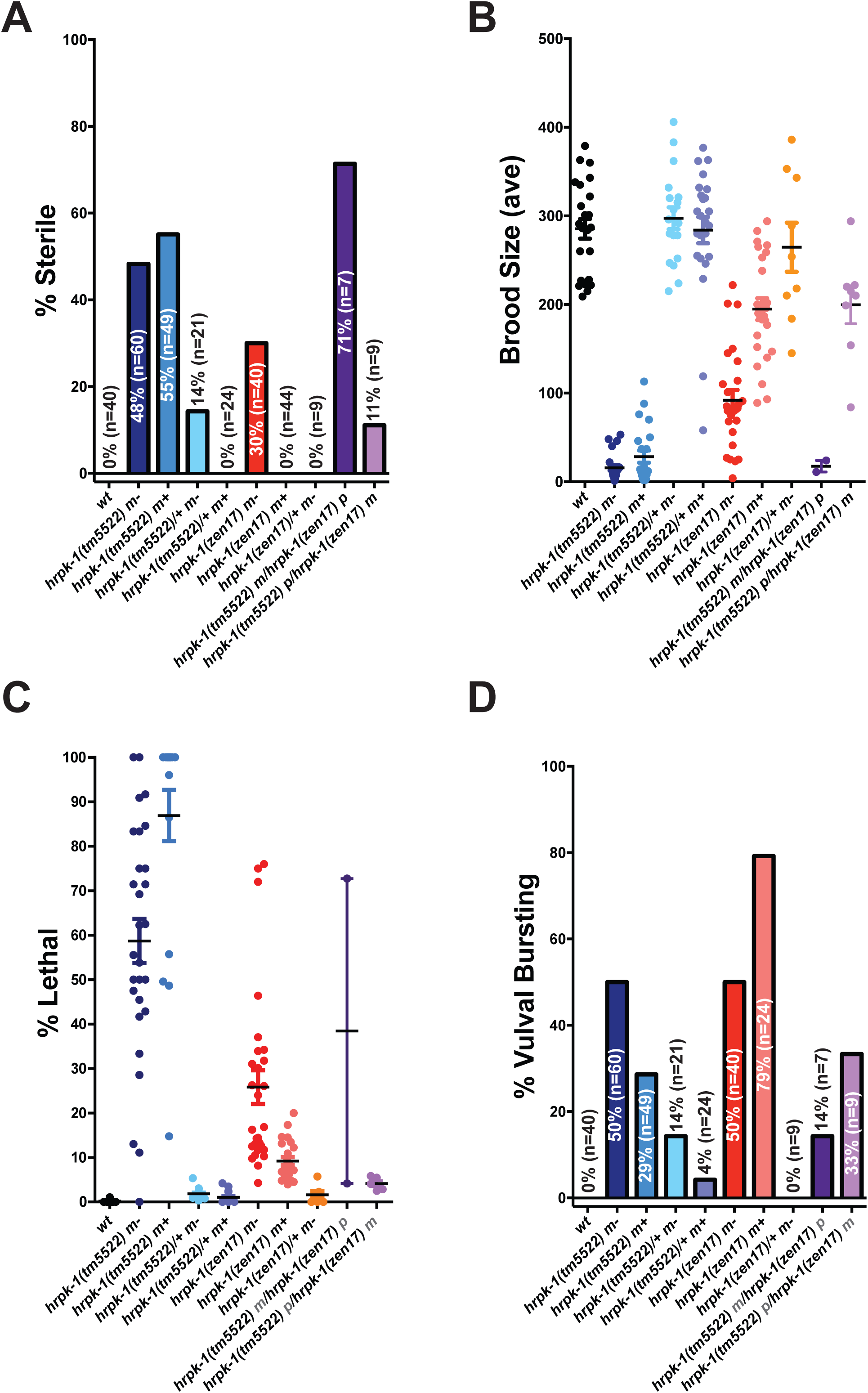
*hrpk-1(tm5522)* is weakly semi-dominant and *hrpk-1* is required maternally. Genetic analyses of *hrpk-1(zen17)* and *hrpk-1(tm5522)* alleles reveal that *hrpk-1* activity has a maternal component for some developmental processes such as fertility (**A**), brood size (**B**), and embryonic viability (**C**), but not for vulval integrity in day 3 or older adults (**D**). *hrpk-1(tm5522)* appears to be weakly semi-dominant as evidenced by the presence of defects observed in *hrpk-1(tm5522)/+* and *hrpk-1(tm5522)/hrpk-1(zen17)* animals (**A-D**). Genotype of the score animals is shown. m-indicates that scored animals came from homozygous mutant mothers, m+ indicates that scored animals were progeny of wild type mothers. *hrpk-1(tm5522)m/hrpk-1(zen17)p* animals came from a cross between *hrpk-1(tm5522)* mothers and *hrpk-1(zen17)* fathers. *hrpk-1(tm5522)p/hrpk-1(zen17)m* animals came from *hrpk-1(zen17)* mothers and *hrpk-1(tm5522)* fathers.

**S3 Fig.**
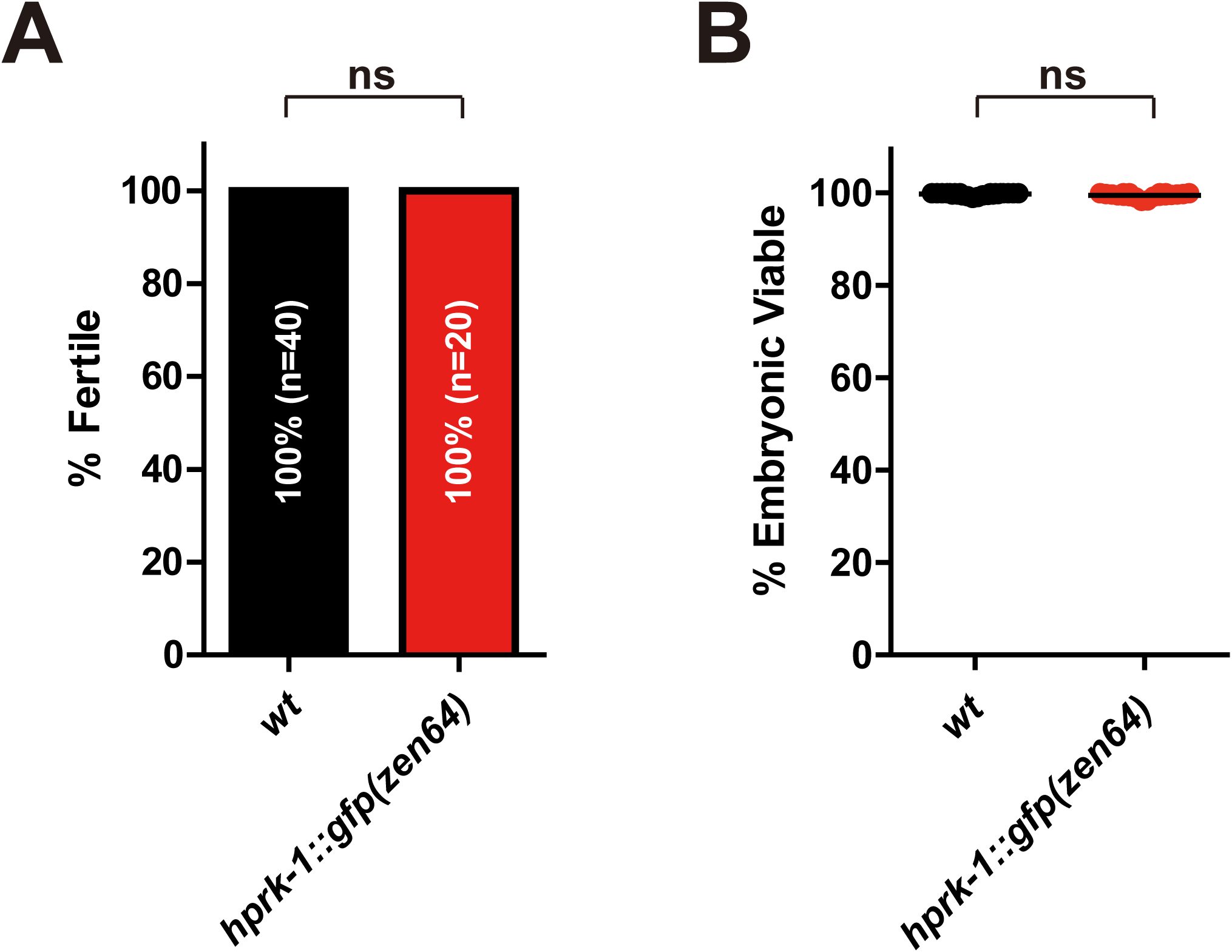
GFP-tagged endogenous HRPK-1 retains its activity. C-terminal HRPK-1 GFP tag does not affect animal fertility (**A**) or embryonic viability (**B**).

**Table S1.**
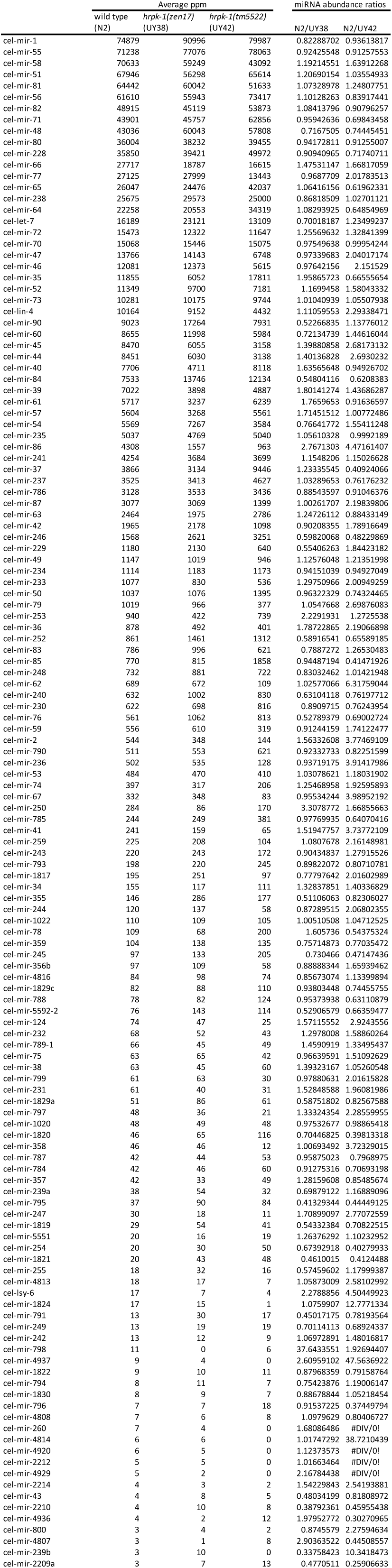
miRNA abundances in *hrpk-1* mutants compared to wild type.

**Table S2.** Mature miRNAs whose abundance was changed 2-fold or more in *hrpk-1* mutants compared to wild type. miRNAs with ≥2-fold decreased abundance are highlighted in gray. miRNAs whose abundace increased in *hrpk-1* mutants ≥2-fold are highlighted in blue.

|  | wild type<br>(N2) | *hrpk-1*(zen17)<br>(UY38) | *hrpk-1*(tm5522)<br>(UY42) | N2/UY38<br>Ratio | N2/UY42<br>Ratio |
| --- | --- | --- | --- | --- | --- |
| cel-mir-798 | 11 | 0 | 6 | 37.643 | 1.927 |
| cel-mir-250 | 284 | 86 | 170 | 3.308 | 1.669 |
| cel-mir-86 | 4308 | 1557 | 963 | 2.767 | 4.472 |
| cel-lsy-6 | 17 | 7 | 4 | 2.279 | 4.504 |
| cel-mir-253 | 940 | 422 | 739 | 2.229 | 1.273 |
| cel-mir-36 | 878 | 492 | 401 | 1.787 | 2.191 |
| cel-mir-247 | 30 | 18 | 11 | 1.709 | 2.771 |
| cel-mir-124 | 74 | 47 | 25 | 1.571 | 2.924 |
| cel-mir-2 | 544 | 348 | 144 | 1.563 | 3.775 |
| cel-mir-41 | 241 | 159 | 65 | 1.519 | 3.738 |
| cel-mir-44 | 8451 | 6030 | 3138 | 1.401 | 2.693 |
| cel-mir-45 | 8470 | 6055 | 3158 | 1.399 | 2.682 |
| cel-mir-797 | 48 | 36 | 21 | 1.333 | 2.286 |
| cel-mir-233 | 1077 | 830 | 536 | 1.298 | 2.009 |
| cel-lin-4 | 10164 | 9152 | 4432 | 1.111 | 2.293 |
| cel-mir-259 | 225 | 208 | 104 | 1.081 | 2.161 |
| cel-mir-1824 | 17 | 15 | 1 | 1.076 | 12.777 |
| cel-mir-4813 | 18 | 17 | 7 | 1.059 | 2.581 |
| cel-mir-79 | 1019 | 966 | 377 | 1.055 | 2.699 |
| cel-mir-62 | 689 | 672 | 109 | 1.026 | 6.318 |
| cel-mir-47 | 409 | 405 | 145 | 1.010 | 2.817 |
| cel-mir-358 | 46 | 46 | 12 | 1.007 | 3.723 |
| cel-mir-87 | 3077 | 3069 | 1399 | 1.003 | 2.198 |
| cel-mir-799 | 61 | 63 | 30 | 0.979 | 2.016 |
| cel-mir-47 | 13766 | 14143 | 6748 | 0.973 | 2.040 |
| cel-mir-77 | 27125 | 27999 | 13443 | 0.969 | 2.018 |
| cel-mir-67 | 332 | 348 | 83 | 0.955 | 3.990 |
| cel-mir-236 | 502 | 535 | 128 | 0.937 | 3.914 |
| cel-mir-244 | 120 | 137 | 58 | 0.873 | 2.068 |
| cel-mir-1817 | 195 | 251 | 97 | 0.778 | 2.016 |
| cel-mir-245 | 97 | 133 | 205 | 0.730 | 0.471 |
| cel-mir-1820 | 46 | 65 | 116 | 0.704 | 0.398 |
| cel-mir-254 | 20 | 30 | 50 | 0.674 | 0.403 |
| cel-mir-246 | 1568 | 2621 | 3251 | 0.598 | 0.482 |
| cel-mir-1821 | 20 | 43 | 48 | 0.461 | 0.412 |
| cel-mir-791 | 13 | 30 | 17 | 0.450 | 0.782 |
| cel-mir-795 | 37 | 90 | 84 | 0.413 | 0.444 |
| cel-mir-37 | 3866 | 3134 | 9446 | 1.233 | 0.409 |

